# A Spatial Attention Guided Deep Learning System for Prediction of Pathological Complete Response Using Breast Cancer Histopathology Images

**DOI:** 10.1101/2022.05.25.493468

**Authors:** Hongyi Duanmu, Shristi Bhattarai, Hongxiao Li, Shi Zhan, Fusheng Wang, George Teodoro, Keerthi Gogineni, Preeti Subhedar, Umay Kiraz, Emiel A.M. Janssen, Ritu Aneja, Jun Kong

## Abstract

Predicting pathological complete response (pCR) to neoadjuvant chemotherapy (NAC) in triple-negative breast cancer (TNBC) patients accurately is direly needed for clinical decision making. pCR is also regarded as a strong predictor of overall survival. In this work, we propose a deep learning system to predict pCR to NAC based on serial pathology images stained with hematoxylin and eosin (H&E) and two immunohistochemical biomarkers (Ki67 and PHH3). To support human prior domain knowledge based guidance and enhance interpretability of the deep learning system, we introduce a human knowledge derived spatial attention mechanism to inform deep learning models of informative tissue areas of interest. For each patient, three serial breast tumor tissue sections from biopsy blocks were sectioned, stained in three different stains, and integrated. The resulting comprehensive attention information from the image triplets is used to guide our prediction system for prognostic tissue regions. The experimental dataset consists of 26,419 pathology image patches of 1,000 × 1,000 pixels from 73 TNBC patients treated with NAC. Image patches from randomly selected 43 patients are used as a training dataset and images patches from the rest 30 are used as a testing dataset. By the maximum voting from patch-level results, our proposed model achieves a 93% patient-level accuracy, outperforming baselines and other state-of-the-art systems, suggesting its high potential for clinical decision making. The codes, the documentation, and example data are available on an open source at: https://github.com/jkonglab/PCR_Prediction_Serial_WSIs_biomarkers

## Introduction

Triple-negative breast cancers (TNBCs) are an aggressive breast cancer subtype with dismal prognosis, high recurrence, and death rates (1). In addition, this subtype has high intratumoral heterogeneity and early stage TNBC tumors lack predictive biomarkers (2). Treatment options for TNBC are mostly limited to chemotherapy and in some instances immunotherapy. Pathological Complete Response (pCR), defined as the disappearance of invasive tumors after neoadjuvant chemotherapy (NAC), is widely used in breast cancer treatment planning (3). Literature suggests that pCR presents a strong positive relationship with the overall survival and disease-free survival (4). However, pCR after NAC treatment is only observed in a limited subset (30-40%) of TNBC patients (5). With an accurate pCR prediction system, those patients predicted as non-or limited responders to NAC treatment could be directed towards clinical trial options that offer alternatives to conventional cytotoxic therapy. Therefore, accurately predicting pCR prior to delivery of neoadjuvant treatment would be critical to personalize treatment.

However, an accurate pCR prediction, prior to chemotherapy, is quite challenging. Despite numerous prior studies using different data modalities such as CT/PET (6, 7), MRI (8, 9), and mammography (10, 11), the prediction performance remains insufficient for clinical deployment. Compared to these data types, hematoxylin and eosin (H&E) stained and IHC-stained whole slide images (WSIs) of tissue biopsies are poorly explored for the prediction of NAC response. WSIs can serve as a decisive image modality that is already routinely used by pathologists for clinical diagnosis and prognosis in clinical settings. While there are few published studies on pCR prediction with pathology images (12–14), the overall prediction performances are limited due to the use of a single information source, e.g., images from a single stain. With the rapid development of deep learning techniques, especially convolutional neural networks (CNNs), recent years have witnessed a significant advance not only in natural image processing research (15), but also in a wide range of biomedical image analysis tasks (16), such as abnormal region detection (17), cell segmentation (18), and survival prediction (19). Despite the great success in medical image analysis tasks, CNNs still present two noteworthy drawbacks: 1) The original CNN design does not make it easy to support multi-modal data processing or multi-modal image analysis. In many medical prognostic tasks, it is often necessary to analyze information from multiple data sources or image types before a final decision can be reached. Although this problem has recently drawn attention from the research community, there is a lack of methods that can be widely proven effective yet (20); 2) The convolution operations in CNNs treat every image pixel equally. However, tissue regions in pathology images do not present an equal amount of prognostic information for the prediction. Paying attention to areas of low or no prognostic values could lead to incorrect or inaccurate disease assessment results (21).

We present a unique solution to address these two problems. In addition to the predictive value from H&E images, another complementary source of information critical for pCR prediction enhancement is Immunohistochemical (IHC) images with informative biomarkers such as Ki67 for cell proliferation and Phosphohistone H3 (PHH3) for mitotic activity (22). In current diagnostic practice, pathologists quantify proliferating cell population within tumors by Ki67 biomarker. Ki67 positivity is also known for its relative responsiveness to chemotherapy (23). The PHH3 mitosis signal characterizes a sub-phase of the entire proliferative cell cycle. Thus, we developed a deep learning pCR prediction system with serial histopathology images including H&E stain, Ki67, and PHH3 IHC biomarkers. Such a prediction system jointly analyzes tumor cells, proliferation, and mitosis biomarkers in a single tissue space that can better capture the profile of cycling proliferating tumor cells and the inherent mitotic propensity in TNBCs. Although CNNs present certain learning abilities for making proper spatial attention on the whole images, the overall robustness, accuracy, and interpretability can be further enhanced with prior human-knowledge (24). We thus propose a deep learning system mimicking human review procedures in clinical settings by creating a spatial attention mechanism specifically on proliferation (Ki67) and mitosis (PHH3) positive tumor cells.

The main contributions of this work are four-fold. First, we developed a deep learning architecture for an enhanced pCR prediction with integrated serial histopathology images stained using H&E stain and two IHC biomarkers (i.e. Ki67 and PHH3). Second, a human knowledge derived spatial attention mechanism is proposed to address the uniform spatial attention limitation by the convolution operations in CNNs. Specifically, the spatial attention mechanism in this work concentrates on regions enriched with proliferative (Ki67^+^) and mitotic (PHH3^+^) tumor cells. Third, the intermediate results from this spatial attention mechanism can also help pathologists identify areas of high prognostic value and improve deep learning interpretability. Finally, the developed spatial attention mechanism can be readily transferred to other customized spatial attention modes to support various tasks in different application scenarios.

## Methods

### System overview

We present the overall architecture of our prediction system in Fig. 1. As we generate whole slide images of the three serially-cut full-face sections stained for H&E and two IHC biomarkers (Ki67 and PHH3), our next processing step is to spatially register these three serial images using a hierarchical based approach, first starting with a low image resolution (25). The resulting learned transformations are next mapped to a high image resolution for retaining the image details. In the top left “Image Registration” panel, registered sample image regions in H&E stain, and of two IHC biomarkers (i.e. Ki67 and PHH3) of size 8,000 × 8,000 and their zoom-in views (1,000 × 1,000 pixels) are presented in the left and right column, respectively. Next, tumor cells in H&E images are detected by a Mask-RCNN based model, while positive biomarker areas (brown) in IHC images are identified by a color deconvolution-based biomarker detector. In the next step, a spatial attention mechanism is introduced to direct the system’s attention to areas enriched with tumor cells of positive Ki67 and PHH3 biomarkers, as such tumor regions suggest noteworthy cell proliferation events and mitotic activities. Finally, a ResNet-34 based deep learning model is trained for pCR prediction.

**Fig. 1.**
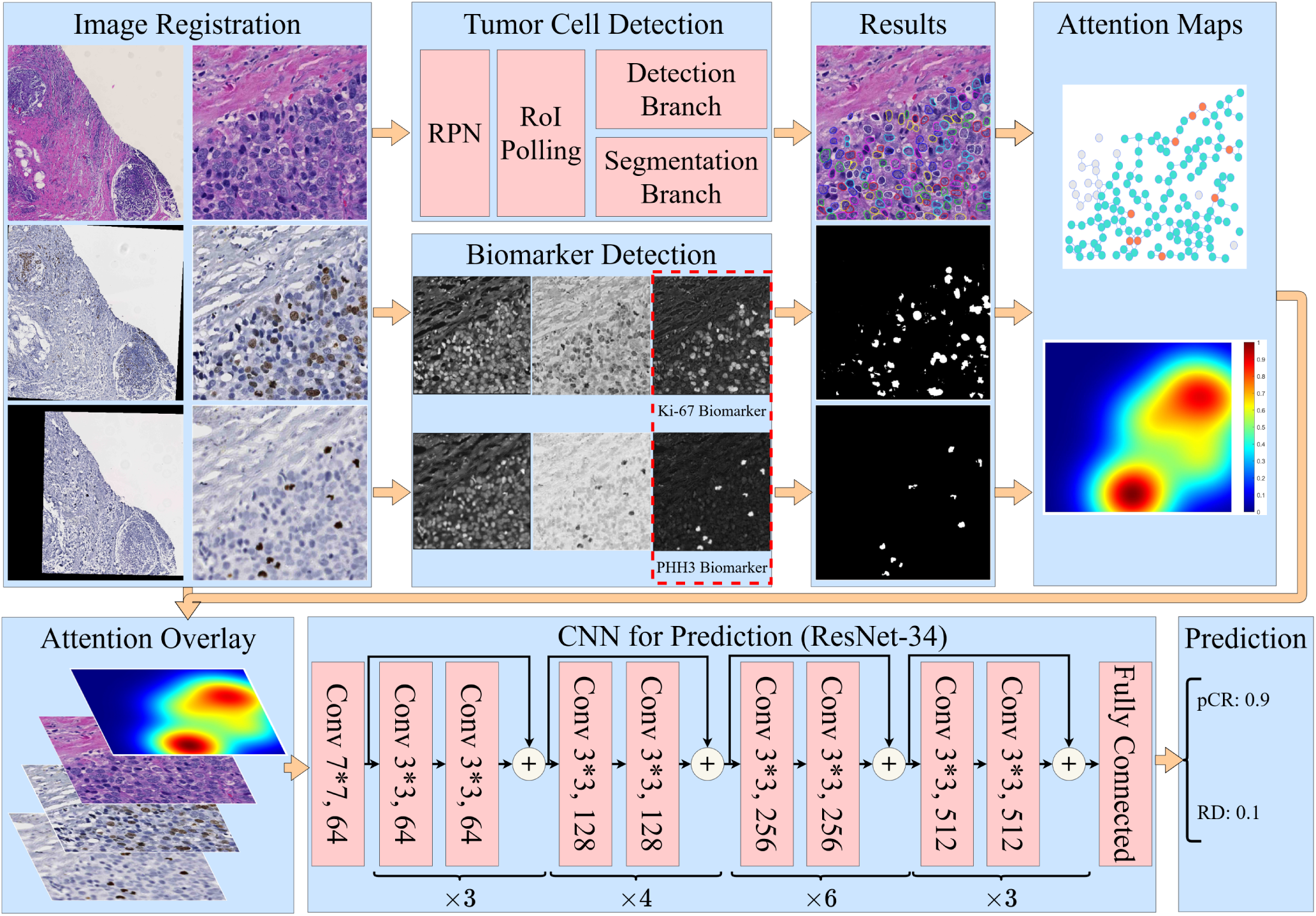
The overall architecture of our proposed system. Serial IHC images with biomarker Ki67 and PHH3 are registered to the H&E slide for each patient. Tumor cells in H&E images are detected, and positive biomarker areas (brown) in IHC images are identified. A spatial attention mechanism is introduced to direct system attention to areas enriched with tumor cells of positive Ki67 and PHH3 biomarkers. Finally, a ResNet-34 based deep learning model is trained for pCR prediction.

### Image registration

To enable the integrated use of serial histopathology images stained using H&E, and two IHC biomarkers, we register such image triplets by a dynamic registration method (25). As each whole slide histopathology image can include giga-pixels by scale, it is technically infeasible to complete whole slide image registration all at once. As a result, the spatial transformations between image pairs are first estimated at a low image resolution. In practice, the highest image resolution that machine memory can accommodate is selected as the low image resolution. With our dataset and machine specifications, the low image level is downscaled by 16 times from the full image resolution. The optimal spatial transformations learned with a rigid and non-rigid registration at the low image resolution are next mapped to the full image resolution. The H&E slide serves as the reference and two IHC images are mapped to the H&E slide. At the high image resolution, the tissue area in the reference H&E slide is divided by 8,000 × 8,000 image regions. Based on the mapped spatial transformations, the corresponding serial Ki67 and PHH3 image regions are aligned to the H&E reference image region after the linear interpolation. To make these mapped IHC image regions match better to the reference, we apply another round of rigid registration for final matched image triplets.

### Cell detection and biomarker detection

As there is a large number of small-scaled tumor cells in histopathology images, Mask-RCNN, a classical two-stage detection system known for its localization accuracy for small object detection, is used as our tumor cell detector (26). Specifically, we customize the anchor scales and ratios in the Mask-RCNN based on tumor cell size in our dataset. Mask-RCNN, which is derived from the Faster-RCNN, has a Region Proposal Network (RPN) for proposing candidate regions containing objects of interest, a Region of Interest (ROI) pooling component for resizing candidate regions, a detection module for classification and the bounding box refinement, and a segmentation head for segmenting proposed regions. Note that our tumor cell detector is trained before the pCR prediction system with independent training datasets. To locate IHC biomarkers in the registered IHC serial images, we constructed an analysis pipeline with the color space conversion and color deconvolution (27). One pre-defined color conversion matrix is applied to all pixels for converting the RGB to the HaematoxylinEosin-DAB (HED) color space. The resulting biomarker positive areas can be recognized by the DAB signal strength. Pixels presenting strong DAB signals are connected spatially and unduly small connected components are removed to enhance the biomarker detection results.

### Spatial attention map generation

We design a domain knowledge derived spatial attention mechanism to guide the deep learning system for specific prognostic areas that pathologists would investigate during the human reviewing process. Rather than inspecting all tumor cells, pathologists usually focus their attention to hot spots enriched with tumor cells that are actively proliferating as suggested by the Ki67 positivity. To identify such informative tissue regions of predictive value to the pCR prediction, we leverage the proliferation and mitosis information from co-registered Ki67 and PHH3 IHC serial images. After identification of tumor cells from the H&E slide and biomarker positive areas, as illustrated in the “Results” panel in Fig. 1. we next detect Ki67^+^ and PHH3^+^ tumor cells. Based on our dataset, the range of tumor cell size is about 60 *−* 100 pixels by the equivalent cell diameter. Therefore, we label a tumor cell to be Ki67 or PHH3 positive if the distance between its center and the nearest Ki67^+^ or PHH3^+^ area is smaller than 100 pixels.

After all tumor cells are labeled by Ki67 and PHH3 biomarker, the prior knowledge derived spatial attention map is produced by the Kernel Density Estimation (KDE) algorithm (28, 29). KDE is a non-parametric way to estimate the probability density function on finite data samples. In this work, the Gaussian kernel function is placed on each tumor cell of positive biomarker and all such responses are summed and normalized to form the smooth density estimation. Some representative attention maps are presented in Fig 2. The kernel bandwidth is set to 150 pixels, which is slightly larger than a tumor cell size to ensure the tumor cell neighborhood coverage. When such spatial attention is generated by the ratio of PHH3^+^ to Ki67^+^ tumor cell number (i.e. Fig. 2(d)), only a PHH3^+^ tumor cell with such a ratio larger than 0.2 within a 500-pixel neighborhood is considered as a valid data sample for the KDE analysis. The kernel bandwidth is set to 500 pixels in that case to capture areas presenting high ratio values. The performances of all attention modes are compared in our study.

**Fig. 2.**
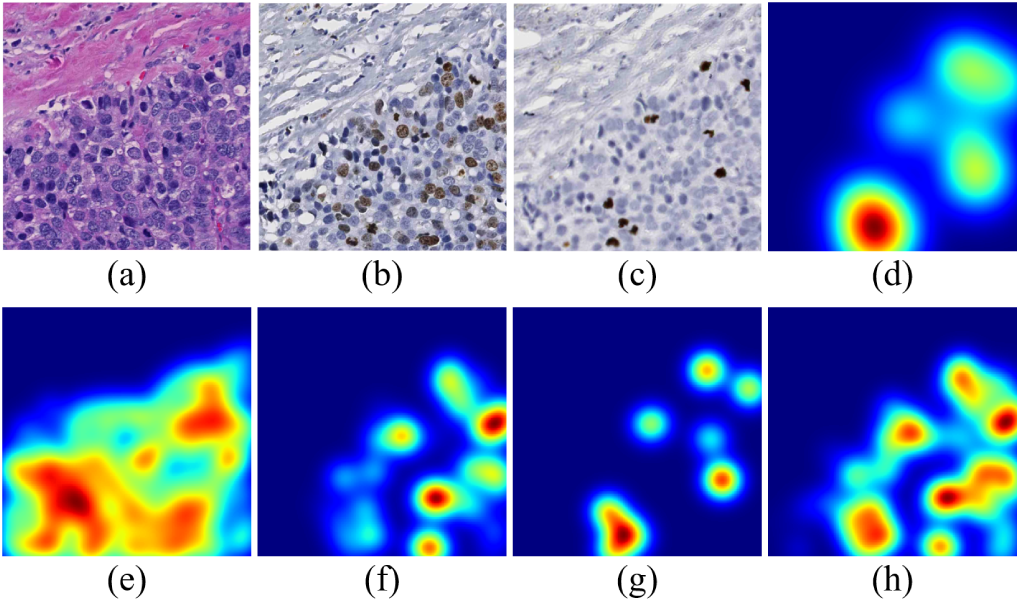
Spatial attention maps of different attention modes for a representative tissue region. (a)-(c): The same small tissue area from co-registered H&E, Ki67, and PHH3 pathology image, respectively; (d): The attention map generated by the ratio of PHH3^+^ to Ki67^+^ tumor cell number; (e)-(g): KDE density estimation results by all tumor cells, Ki67^+^ tumor cells, and PHH3^+^ tumor cells, respectively; (h) KDE density estimation results by the sum of Ki67^+^ and PHH3^+^ tumor cell numbers.

### pCR prediction

The resulting spatial attention maps are multiplied with original pathology images in a pixel-wise manner. The resulting attention images are provided to the prediction module that uses ResNet-34 as the backbone (30). Given the computation burden, data scale, and model complexity, ResNet-34 is a suitable backbone for our study. After the first convolutional layer with a kernel size of 7, four main blocks having different channel numbers (i.e. 64, 128, 256, and 512, respectively) are proceeded with for further analysis. For better feature extraction, residual sub-blocks in these main blocks are repeated for 3, 4, 6, and 3 times, respectively. In each residual sub-block, two convolutional layers with a kernel size of 3 are included. Additionally, one shortcut connection characteristic of the ResNet is deployed. All convolutional layers are followed by an activation function layer and a batch normalization layer. Finally, fully connected layers are deployed to make the pCR prediction after the global average pooling.

## Results

### Data description and training configurations

A dataset for tumor cell detection independent of the dataset for pCR prediction is used to train the tumor cell detector. The tumor cell training dataset consists of 868 40x H&E histopathology image regions of size 8,000 × 8,000 pixels. In total, 53,314 tumor cells are manually annotated. Due to the large size of these image regions, they are partitioned to non-overlapped 1,000 × 1,000 image patches that are next resized to 512 × 512 pixels to fit the deep learning tumor cell detector. The tumor cell detector is trained for 200 epochs by the cross-entropy loss with an NVIDIA V100 GPU. In our study, Stochastic Gradient Descent (SGD) is used as the optimizer. Nesterov momentum is activated and the learning rate is set as 0.001.

A separate dataset of serial histopathology images in H&E and of Ki67 and PHH3 IHC biomarkers is used for pCR prediction training. A total of 73 NAC treated TNBC cases are collected before neoadjuvant therapy from Emory University Healthcare. Formalin-fixed and paraffin-embedded serial tissue biopsies from each patient before neoadjuvant therapy are H&E stained and immunohistochemically stained for Ki67 and PHH3. The resulting slides are digitally scanned and co-registered. pCR cases are those biopsy pretreatment specimens whose post-NAC surgical specimens had no residual invasive carcinoma in both the breast tissue and regional lymph nodes. Our analysis is based entirely on the evaluation of the pre-treatment biopsy; we did not analyze the post-NAC treatment specimens. If a patient is not pCR, it is considered a residual disease (RD) case. Based on these criteria, 43 and 30 patients are labeled with pCR and RD after NAC, respectively.

A data summary of our dataset for prediction is presented in Table 1. Tissue regions of all 73 patients result in 969 40x histopathology image regions of size 8,000 × 8,000 pixels. The training dataset for pCR prediction includes randomly selected 26 pCR and 17 RD patients with 553 tissue region images of size 8,000 × 8,000 pixels. Meanwhile, the testing dataset includes the other 17 PCR and 13 RD cases with 416 pathology image regions of size 8,000 × 8,000 in pixels. Each such image region is partitioned into non-overlapped 1,000 × 1,000 image patches that are next resized to 512 × 512 images for deep learning model analysis. Images with less than 20 tumor cells are excluded from the dataset to ensure data for analysis are properly represented. A total of 16,539 and 9,880 pathology images patches are included in the training and testing dataset, respectively. As the resulting positive-to-negative ratio is around 1.5, our dataset is generally balanced.

**Table 1.** Summary of the dataset for pCR prediction. Image numbers and the ratios of positive over negative cases in patient, region, and patch levels are provided respectively.

|  |  | pCR | RD | Total | Pos/Neg |
| --- | --- | --- | --- | --- | --- |
| Training | Patient level | 26 | 17 | 43 | 1.53 |
|  | Region level | 313 | 240 | 553 | 1.30 |
|  | Patch level | 9,321 | 7,218 | 16,539 | 1.29 |
| Testing | Patient level | 17 | 13 | 30 | 1.31 |
|  | Region level | 266 | 150 | 416 | 1.77 |
|  | Patch level | 6,249 | 3,631 | 9,880 | 1.72 |
| Total | Patient level | 43 | 30 | 73 | 1.43 |
|  | Region level | 579 | 390 | 969 | 1.48 |
|  | Patch level | 15,570 | 10,849 | 26,419 | 1.44 |

### Model evaluation

We implemented and compared six different models to evaluate the feasibility and efficacy of the proposed human knowledge guided spatial attention mechanism targeting Ki67^+^ and PHH3^+^ tumor cells. They are pCR prediction systems 1) with no attention mechanism; 2) with an attention mechanism on all tumor cells; 3) with an attention mechanism on Ki67^+^ tumor cells; 4) with an attention mechanism on PHH3^+^ tumor cells; 5) with an attention mechanism on a high *γ*% where *γ*% is defined as the percentage of Ki67^+^ tumor cells presenting PHH3^+^ signals; and 6) with an attention mechanism on both Ki67^+^ and PHH3^+^ tumor cells. Evaluation metrics, including Accuracy, Area Under Curve (AUC), Precision, and Recall, are used for quantitative evaluation.

The detailed quantitative performance results are presented in Table 2. Note the pCR prediction decisions at the region and patient level are made by the maximum voting from the patch and region level, respectively. The prediction system without any spatial attention mechanism presents the least competitive performance, with 0.700, 0.702, and 0.733 Accuracy at the patch, region, and patient level, respectively. With spatial attention on tumor cells, the model Accuracy is improved to 0.783, 0.800, and 0.800 at the patch, region, and patient level, respectively. Noticeable improvements are also suggested by other metrics. Additionally, the spatial attention on either Ki67^+^ or PHH3^+^ tumor cells is further beneficial to an enhanced pCR prediction, achieving AUCs by 0.825 and 0.802 at the patch level, and accuracy by 0.860 and 0.836 at the region level, respectively. Additionally, both sources of attention achieve 0.833 Accuracy at the patient level. Noteworthy models in the last two columns, i.e. Tumor (*γ*%), and Tumor (Ki67^+^ PHH3^+^), boost the performance even further. The model with the attention on *γ*% is better than the previous four models by 6.42%, 8.14% and 7.18% for Accuracy at the patch, region, and patient level, respectively. By contrast, our best model with the attention on both Ki67^+^ and PHH3^+^ tumor cells is better than the previous four models by 8.24%, 9.30%, and 12.00% for Accuracy at the patch, region, and patient level, respectively.

**Table 2.** Quantitative performance comparison across all proposed models by Accuracy, AUC, Precision, and Recall at the image patch, region, and patient level.

| | | w/o attention | Attention on tumor | Tumor (Ki67<sup>+</sup>) | Tumor (PHH3<sup>+</sup>) | Tumor ($\gamma\%$) | Tumor (Ki67<sup>+</sup> ∪ PHH3<sup>+</sup>) |
| --- | --- | --- | --- | --- | --- | --- | --- |
| Patch level | Accuracy | 0.700 | 0.783 | 0.825 | 0.802 | 0.878 | **0.893** |
|  | AUC | 0.710 | 0.812 | 0.884 | 0.838 | 0.946 | **0.962** |
|  | Precision | 0.722 | 0.784 | 0.828 | 0.801 | **0.908** | 0.897 |
|  | Recall | 0.854 | 0.907 | 0.912 | 0.914 | 0.898 | **0.939** |
| Region level | Accuracy | 0.702 | 0.800 | 0.860 | 0.836 | 0.930 | **0.940** |
| Patient level | Accuracy | 0.733 | 0.800 | 0.833 | 0.833 | 0.900 | **0.933** |

By performance comparison, the prediction system with spatial attention on *γ*% is slightly worse than the system with spatial attention on both Ki67^+^ and PHH3^+^ tumor cells by all metrics but Precision. As Precision and Recall are related to type I and II error, the system with spatial attention on *γ*% presents an overall inferior performance to our system with attention on Ki67^+^ and PHH3^+^ tumor cells if both metrics are combined. As the latter system learns attention from Ki67^+^ and PHH^+^ cells in a freestyle manner, it can generalize the attention information better than the system learning attention on pre-defined *γ*% measure. The Receiver Operator Characteristic (ROC) curve demonstrated in Fig. 3(a) confirms this observation.

**Fig. 3.**
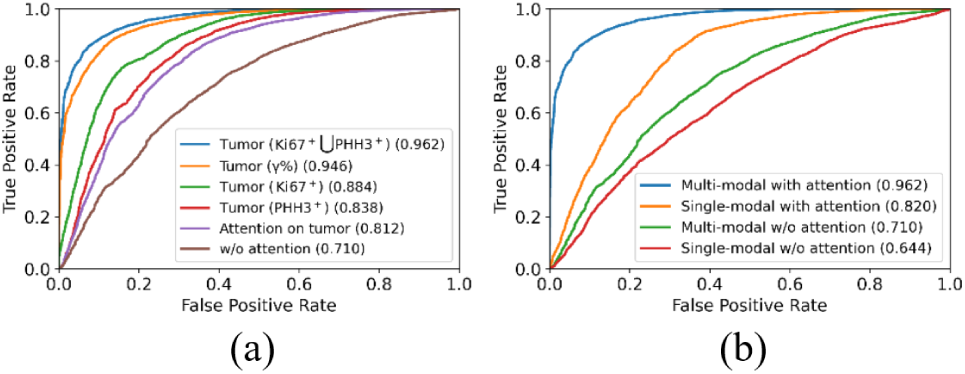
Comparison of ROC curves of all evaluated models (a) using distinct attention information, and (b) for ablation studies.

### Ablation study

There are two salient features that make our proposed model unique. It integrates serial histopathology H&E image with Ki67 and PHH3 (IHC) biomarker stained images and includes a domain knowledge guided spatial attention learning mechanism. We design an ablation study to evaluate these two components. Table 3 presents ablation study results of four models 1) processing only H&E slides without any spatial attention mechanism (i.e. Model 1), 2) processing multi-modal serial slides without any spatial attention mechanism (i.e. Model 2), 3) processing only H&E slides with our best spatial attention mechanism (i.e. Model 3), and 4) processing multi-modal serial slides with our best spatial attention mechanism (i.e. Model 4), respectively.

**Table 3.** Quantitative performance comparison among all ablated models for the ablation study.

|  |  | Single-modal w/o attention | Multi-modal w/o attention | Single-modal with attention | Multi-modal with attention |
| --- | --- | --- | --- | --- | --- |
| Patch level | Accuracy | 0.665 | 0.700 | 0.800 | 0.893 |
|  | AUC | 0.644 | 0.710 | 0.820 | 0.961 |
|  | Precision | 0.679 | 0.722 | 0.802 | 0.897 |
|  | Recall | 0.894 | 0.854 | 0.908 | 0.939 |
| Region level | Accuracy | 0.673 | 0.702 | 0.815 | 0.940 |
| Patient level | Accuracy | 0.600 | 0.733 | 0.800 | 0.933 |

Without attention information from IHC biomarkers, the baseline system (i.e. Model 1) yields 0.665, 0.673, and 0.600 Accuracy at the patch, region, and patient level, respectively. With the inclusion of serial pathology IHC images for prediction in Model 2, the prediction performance is improved to 0.700, 0.702, and 0.733 Accuracy at the patch, region, and patient level, respectively. Similar system performance enhancement patterns are also observed between Model 3 and 4. Additionally, the high Recall and low Precision in both systems with only H&E stained images (i.e. Model 1 and 3) indicate that these systems tend to produce false positives by predicting a case more preferably as pCR. Without IHC Ki67 and PHH3 biomarker information support, H&E pathology image based prediction, especially without any spatial attention, only achieves limited success.

Comparing models without (i.e. Model 1 and 2) and with prior domain knowledge based attention (i.e. Model 3 and 4), we notice that our proposed spatial attention mechanism drastically improves pCR prediction by 20.30% between Model 1 and 3, and 27.57% between Model 2 and 4 at the patch level, respectively. The improvement percentages are 21.10% and 33.90% at the region level, and 33.33% and 27.29% at the patient level. All these results suggest a clear efficacy of our proposed domain knowledge guided spatial attention mechanism for pCR prediction. Additionally, ROC curves plotted in Fig. 3(b) suggest the same conclusions.

### Visualization

To demonstrate the critical role our proposed domain knowledge attention plays for pCR prediction, we present four typical sample cases in Fig. 4. The first three columns from left to right are four registered H&E, Ki67, and PHH3 histopathology images regions, respectively. The spatial attention maps generated from Ki67^+^ and PHH3^+^ tumor cells are presented in the fourth column. With class activation map (CAM) algorithm (31), we produce the spatial attention maps in the last two columns from prediction models with and without the spatial attention mechanism (i.e. Model 2 and 4). CAM algorithm is designed to visualize deep learning results. It replaces the fully connected layers after the last convolutional layer with one global pooling layer and one new fully connected layer. The resulting system is trained again by only allowing perturbations of newly added model weights. After model training, a heatmap can be produced by a linear combination of weighted feature maps from the last convolutional layer for each testing image. Such heatmaps indicate image attention value for prediction. By attention maps in Fig. 4, the CAM visualization results from the prediction system with the spatial attention are similar to the prior domain knowledge derived spatial attention maps, suggesting a good attention learning outcome. Informative tissue areas for pCR prediction by human prior knowledge are properly analyzed by the prediction system with spatial attention. By contrast, the CAM visualization results from the system without an attention mechanism tend to cover wider tissue regions with a weaker specificity for locating informative regions.

**Fig. 4.**
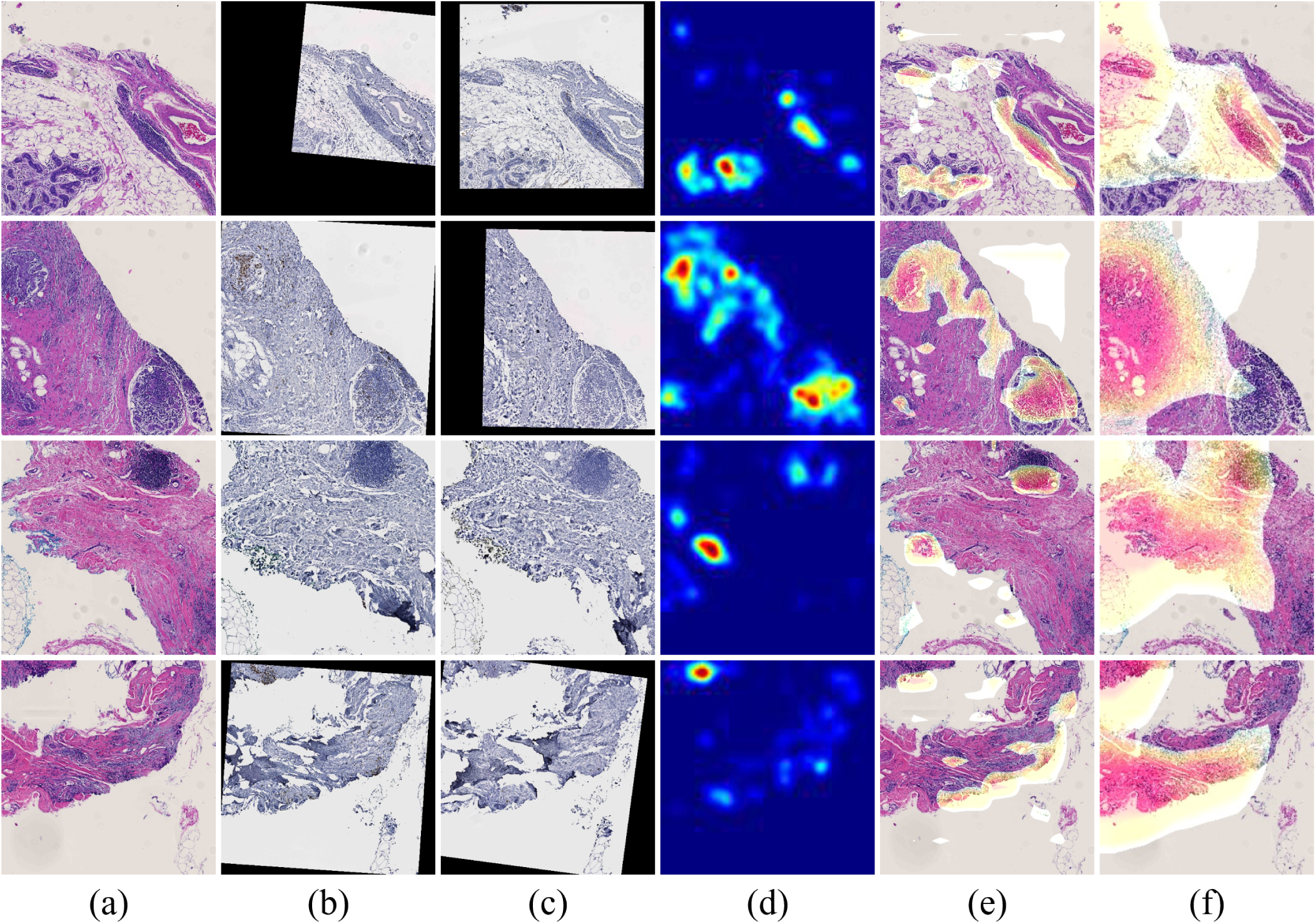
Attention visualization demonstration for typical tissue regions. Representative H&E, Ki67, and PHH3 histopathology image regions are shown in (a)-(c); (d) Spatial attention maps targeting Ki67^+^ and PHH3^+^ tumor cells; CAM visualization results from prediction models (e) with and (f) without the spatial attention mechanism.

### Efficiency

The computational efficiency of a computer-aided clinical supporting system is important as it may limit a system deployment potential in real clinical settings. By a computer cluster equipped with one NVIDIA Tesla V100 GPU and 2 Intel Xeon CPUs with 88 cores, it takes less than 5 seconds to detect tumor cells in a H&E histopathology image region of size 8,000 × 8,000 pixels. It takes about 10 seconds to finish the biomarker detection process. Although the time cost for the spatial attention map generation highly depends on the number of Ki67^+^ and PHH3^+^ tumor cells in an image, the average cost by our dataset is about 10 seconds. The final prediction module is efficient and can complete the prediction of an image input in 2 seconds. When I/O time cost is included, the overall PCR prediction analysis can be completed in 30 seconds per image triplet of 8,000 × 8,000 pixels by size. Equipped with parallel computing and batch operations, such analyses can be sufficiently boosted for clinical support.

## Discussion

### Comparison to other state-of-the-art systems

To our best knowledge, few studies have been reported on the pCR prediction using histopathology images of pretreatment biopsies of patients with TNBCs utilizing machine learning, especially deep learning methods. For example, pCR status has been predicted with manually designed features from tumor infiltrating lymphocytes and their subtypes (12). The number and spatial density of detected tumor infiltrating lymphocytes are provided to the univariate and multivariate logistic regression with other clinical variables including age, tumor size, tumor type, and molecular subtype. The best prediction performance by the odds ratio is 6.44. Similar studies on pCR prediction by logistic regression with cellular information have been reported (13, 32). One study uses cell quantity profiles from pathology images for pCR prediction by conventional machine learning methods, achieving an odds ratio of 4.46 (32). In a follow-up investigation from the same group, the cell density from local neighborhoods in H&E slides is used for prediction, but with a lower odds ratio of 2.92 (13). In another study, pCR is predicted by tumor cell features (e.g. count, size, and circularity) and image based features (e.g. mean pixel intensity and correlation of the gray-level co-occurrence matrix). By the multivariate binary logistic regression, it achieves an accuracy of 0.79 in a dataset with 58 patients (33). All these studies using conventional machine learning methods present a limited prediction success, largely due to limited sources of information and human-defined engineering features insufficient for prediction support.

By contrast, our proposed prediction system overcomes these limitations by a human prior knowledge guided deep learning attention framework with integrative use of serial pathology slides in multiple stains, achieving much better performance with an odds ratio higher than 50. Except for our previous study on this topic (34), only one study using deep learning for this prediction problem is found to our best knowledge (14). The Inception-V3 based CNN architecture is reported to predict pCR from NAC from pre-determined tumor epithelium regions in H&E pathology images, achieving 0.84 by AUC. When tumor epithelium regions from H&E images are combined with information on stromal tumor-infiltrating lymphocytes and tumor subtype, the prediction performance is improved to 0.89 by AUC. By comparison, our system integrating H&E stained pathology images with IHC Ki67, and PHH3 biomarkers from serial slides achieves 0.96 by AUC at the patch level. Encouraged by promising prediction results from our previous work (34), we further introduce human prior knowledge guided spatial attention mechanism and make such embedded attention module readily customizable to other prediction analyses.

### Spatial attention mechanism

Whole slide histopathology images are often large in size with giga-pixels. Not all tissue areas present equal prognostic values for prediction analyses. As a result, it is important to accurately locate image areas with high predictive values for further processing. This work introduces domain knowledge driven spatial attention to guide deep learning model training. In this way, deep learning models can pay more attention to prognostic tissue areas, contributing to an enhanced training outcome. Leveraging the clinical biomarker Ki67 for proliferation and the mitosis activity signal for better tumor cell cycling characterizations (22**?**), we develop a spatial attention mechanism for a deep learning model to target tumor regions enriched with Ki67^+^ and PHH3^+^ tumor cells, effectively reducing the negative learning impact from misleading areas. Different from our prior work (34), we make the spatial attention mechanism separated from the final prediction module. In this way, spatial attention mode can be easily transferred to other attention modes optimal for other studies.

We present one representative patient H&E whole slide image and four sampled tissue regions that include spatial attention by our method in Fig. 5. With our fully automated analysis pipeline, the resulting spatial attention maps can facilitate pathologists to identify prognostic tissue areas of interest in a large whole slide image. In the meanwhile, such an attention mechanism can help improve the interpretation of how a deep learning model works, which is critical to an informed clinical decision support.

**Fig. 5.**
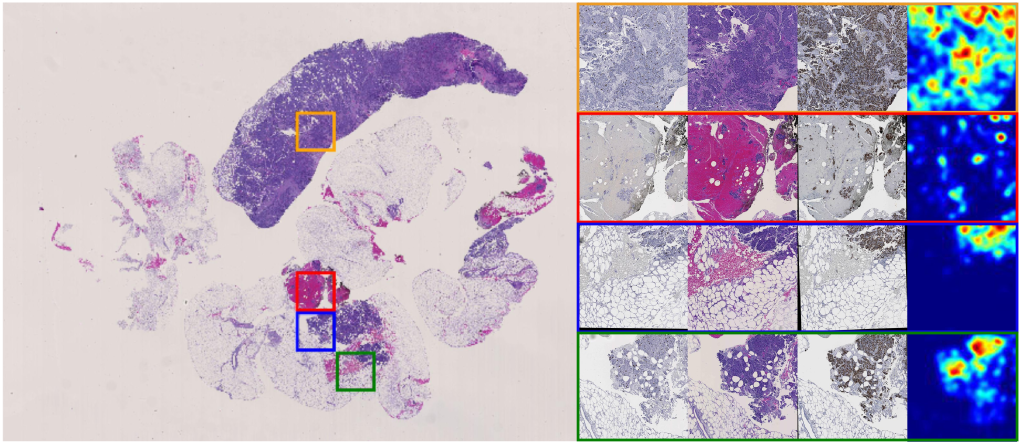
Attention for a typical case is presented. The H&E whole slide image is presented on the left panel. Close up views of four sampled regions in H&E, Ki67, and PHH3 and their attention maps are presented on the right panel.

### IHC biomarker measure

Although proved to be promising by our comparison experiments, the prediction system focusing on PHH3^+^ tumor cells slightly underperforms when it is compared with the model on Ki67^+^ tumor cells overall. Notably, there are fewer positive biomarker areas in PHH3 than those in Ki67 IHC images. With too few tissue areas to focus on, it can result in insufficient system robustness for an accurate prediction. Additionally, the prediction model with the attention on *γ*% is slightly worse than the one with attention on both the Ki67^+^ and PHH3^+^ tumor cells. This can result from two major reasons. First, the measure *γ*% is sensitive to the computation errors from prior analysis steps, including the image registration, cell localization, and biomarker detection. Second, the measure *γ*%, as a manually pre-defined parameter, may not be the best feature representation that can be learned from IHC images. By contrast, the deep learning model can extract the optimal feature patterns to support prediction analysis in principle.

### Model implementation

We design and implement our prediction system by principles of both efficacy and efficiency. Considering the inference speed, portability, and detection performance on small objects, the two-stage detector, Mask R-CNN, is chosen. One-stage detection systems generally present worse detection performance for small and crowded targets of interest (35). By the same criteria, a variant of ResNet architecture, i.e. ResNet-34, is chosen for its balanced efficiency and effectiveness. Moreover, we increase the input image size from 224 × 224 used in the original ResNet-34 to size 512 × 512 to achieve a larger scope of tissue perception but without too much computation burden increase.

### Limitations and future work

There are several directions we plan to investigate in future work. First, we use serial tissue sections capturing both cellular phenotype and molecular biomarker information; thus the resulting dataset scale is limited. Our ongoing efforts are focused on collecting data from more patients for more comprehensive system training and testing. Second, as a key step to enable information integration from serial histopathology images, serial image registration affects all the following analysis steps. It is thus important to further improve the registration accuracy and the system robustness with imperfect inputs. Third, in our current analysis pipeline, the final prediction decision is made by the maximum voting of local image patch level predictions. This decision-making mechanism limits the system from leveraging global tissue information. A global analysis schema is needed to enhance the divide-combine mode.

## Conclusions

In this work, we develop a domain knowledge guided deep learning system to predict pCR to NAC in a TNBC cohort. The developed system simultaneously processes and integrates three serial pathology images capturing complementary tissue phenotype and molecular information critical to pCR prediction. The newly proposed spatial attention mechanism directs deep learning attention to tissue regions with prognostic values by human prior knowledge. The intermediate attention results are visualized to help pathologists locate predictive tissue regions and interpret deep learning predictions. With a well-prepared TNBC patient dataset, our system achieves 93% accuracy for pCR prediction. With a systematic ablation study, we demonstrate the efficacy of an integrative use of tissue phenotype and molecular information from serial tissue slides and the proposed domain knowledge driven spatial attention mechanism. The great prediction performance, highly generalized attention, and strong system interpretability of our prediction system suggest its promising clinical and translational impact on enhancing breast cancer treatment planning.

